# Citrusgreening.org: An open access and integrated systems biology portal for the Huanglongbing (HLB) disease complex

**DOI:** 10.1101/868364

**Authors:** Mirella Flores-Gonzalez, Prashant S Hosmani, Noe Fernandez-Pozo, Marina Mann, Jodi L. Humann, Dorrie Main, Michelle Heck, Susan Brown, Lukas A Mueller, Surya Saha

## Abstract

We have created an open access web portal with pathosystem-wide resources and bioinformatics tools for the host citrus, the vector Asian citrus psyllid (ACP) and multiple pathogens including *Ca*. Liberibacter asiaticus. To the best of our knowledge, this is the first example of a database to use the pathosystem as a holistic framework to understand an insect transmitted plant disease. This endeavor integrates and enables the analysis of data sets generated by the community to study the citrus greening disease complex. Users can submit relevant data sets to enable sharing and allow the community to better analyze their data within an integrated system. The portal contains a variety of tools for omics data. Metabolic pathway databases, CitrusCyc and DiaphorinaCyc provide organism specific pathways that can be used to mine metabolomics, transcriptomics and proteomics results to identify pathways and regulatory mechanism involved in disease response. Psyllid Expression Network (PEN) contains expression profiles of ACP genes from multiple life stages, tissues, conditions and hosts. Citrus Expression Network (CEN) contains public expression data from multiple tissues and conditions for various citrus hosts. All tools like Apollo/JBrowse, Biocyc, Blast, CEN and PEN connect to a central database containing gene models for citrus, ACP and multiple Liberibacter pathogens. The portal also includes electrical penetration graph (EPG) recordings of ACP feeding on citrus, information about citrus rootstock trials and metabolomics data in addition to traditional omics data types with a goal of combining and mining all information related to a pathosystem. The portal includes user-friendly manual curation tools to allow the research community to continuously improve this knowledge base as more experimental research is published. Bulk downloads are available for all genome and annotation datasets from the FTP site (ftp://ftp.citrusgreening.org). The portal can be accessed at https://citrusgreening.org/.

## Introduction

Citrus greening or Huanglongbing (HLB) is a tritrophic disease complex involving citrus, the Asian citrus psyllid (ACP, *Diaphorina citri)* vector and a phloem restricted bacterial pathogen *Ca*. Liberibacter asiaticus (CLas). HLB is considered to be the most devastating of all citrus diseases, and there is currently no adequate control strategy (Wang 2019). In Florida, an estimated 40-70% of all citrus trees are infected and HLB effects include production decline, diminished fruit quality and increased production costs. Further economic losses have been realized in the loss of more than 6,600 jobs in citrus and citrus related sectors since 2012 (Hodges and Spreen, 2012) with an estimated average crop reduction of 40% compared to healthy trees (Singerman, 2015). Citrus greening disease is a devastating disease that is ravaging the citrus industry in the US and world-wide (Wang 2019). Thus, studies are required to understand varietal-specific impacts of citrus greening on tree metabolism and health in different environmental conditions in addition to understanding the role of the insect vector. A major challenge in studying eukaryotic multiple component diseases at a genome-wide scale is the lack of tools for working with non-model systems. This challenge is exacerbated in the case of invasive disease systems such as HLB impacting important food crops where the knowledge about the epidemiology is limited but rapid solutions to reduce the transmission and identify resistance are needed to manage the disease in the field.

Here, we have designed a web portal with information for citrus growers, plant geneticists, vector biologists and plant pathologists. Citrusgreening.org is an open access and public web portal with pathosystem-wide resources and bioinformatics tools for citrus, ACP and multiple pathogens including CLas. This database uses the pathosystem as a holistic framework to understand the biology of disease transmission and response across the host plant, insect vector and bacterial pathogen by integrating datasets generated by the community to study the citrus greening disease complex.

The portal contains a variety of tools for omics data. It provides basic features to query omics data with BLAST databases and visualization with the JBrowse genome browser (Buels et al. 2016) and Apollo annotation tool (Dunn et al. 2019). DiaphorinaCyc and CitrusCyc pathway databases can be used to analyze transcriptomics and proteomics results to identify pathways with differentially regulated genes. Psyllid Expression Network (PEN) contains profiles of ACP genes from multiple life stages, conditions and hosts. Citrus Expression Network (CEN) contains public expression data for citrus from NCBI. The user-friendly manual curation tools will allow the research community to continuously improve this knowledge base as more experimental research is published and we learn more about the interactions within the pathosystem. All tools connect to a central database containing gene models for citrus, ACP and multiple Liberibacter pathogens. The portal can be accessed at https://citrusgreening.org/.

### Gene-centric database

Genome sequencing is typically the first mode of investigation for non-models systems therefore genome sequences form the primary component of the CitrusGreening.org database (Figure 1). The portal contains genomes and annotation for *Citrus clementina* and *Citrus sinensis* among host species besides the vector *Diaphorina citri* and the pathogen *Candidatus* Liberibacter asiaticus psy62. Genomic data is stored in the CHADO relational schema (Mungall et al. 2007). This enables visualization of genomic features along with associated metadata. The genes and other annotated features are searchable from the feature search page (https://citrusgreening.org/search/features). The feature page (Figure 2) is a hub for all the information available for a given gene such as sequence data with links to the BLAST tool for further analysis and annotations with links to external databases like NCBI. The blast databases contain proteins, genomes, coding sequence and transcripts for all reference genomes (Table 1) in addition to other sequence sets of interest (Supplementary Table 1). The feature page and BLAST results link to the Jbrowse genome browser (Buels et al. 2016) to visualize annotations on the genomic backbone. Bulk downloads are available for all genome and annotation datasets from the FTP site (ftp://ftp.citrusgreening.org).

**Table 1.** Genomic resources available at citrusgreeening.org

|  | Genome | Resources | Reference |
| --- | --- | --- | --- |
| Host | *Citrus clementina*, 182 v1 (Clementine mandarin) | BLAST, Jbrowse, Apollo, CitrusCyc pathway database, Expression Network (CEN) | (Wu et al. 2014) |
|  | *Citrus sinensis*, v2 (Sweet orange) | BLAST, Jbrowse, Apollo, CitrusCyc pathway database, Expression Network (CEN) | (Xu et al. 2013) |
|  | *Citrus sinensis*, 154 v1 (Sweet orange) | BLAST, Jbrowse, Apollo, CitrusCyc pathway database | (Wu et al. 2014) |
| Vector | *Diaphorina citri* (Asian citrus psyllid) v3 | BLAST, Jbrowse, Apollo | Unpublished |
|  | *Diaphorina citri* (Asian citrus psyllid) v2 | BLAST, Jbrowse, Apollo, DiaphorinaCyc pathway database, Expression Network (PEN) | Unpublished |
|  | *Diaphorina citri* (Asian citrus psyllid) v1 | BLAST, Jbrowse, Apollo, DiaphorinaCyc pathway database,<br>Expression Network<br>(PEN) | (Saha et al. 2019) |
| Pathogen | *Candidatus Liberibacter asiaticus* psy62<br>(CLas) | BLAST, Jbrowse,<br>Apollo,<br>CitrusgreeningCyc<br>pathway database | (Duan et al.<br>2009) |
|  | *Candidatus Liberibacter asiaticus* gxpsy | BLAST, Jbrowse,<br>Apollo,<br>CitrusgreeningCyc<br>pathway database | (Lin et al.<br>2013) |
|  | *Candidatus Liberibacter asiaticus* Ishi-1 | BLAST, Jbrowse,<br>Apollo,<br>CitrusgreeningCyc<br>pathway database | (Kato et al.<br>2014) |
|  | *Candidatus Liberibacter americanus*<br>São Paulo | BLAST, Jbrowse,<br>Apollo,<br>CitrusgreeningCyc<br>pathway database | (Wulff et al.<br>2014) |
|  | *Candidatus Liberibacter solanacearum*<br>CLso ZC-1 | BLAST, Jbrowse,<br>Apollo,<br>CitrusgreeningCyc<br>pathway database | (Lin et al.<br>2011) |
|  | *Liberibacter crescens* BT-1 | BLAST, Jbrowse,<br>Apollo,<br>CitrusgreeningCyc<br>pathway database | (Leonard et<br>al. 2012) |

**Figure 1:**
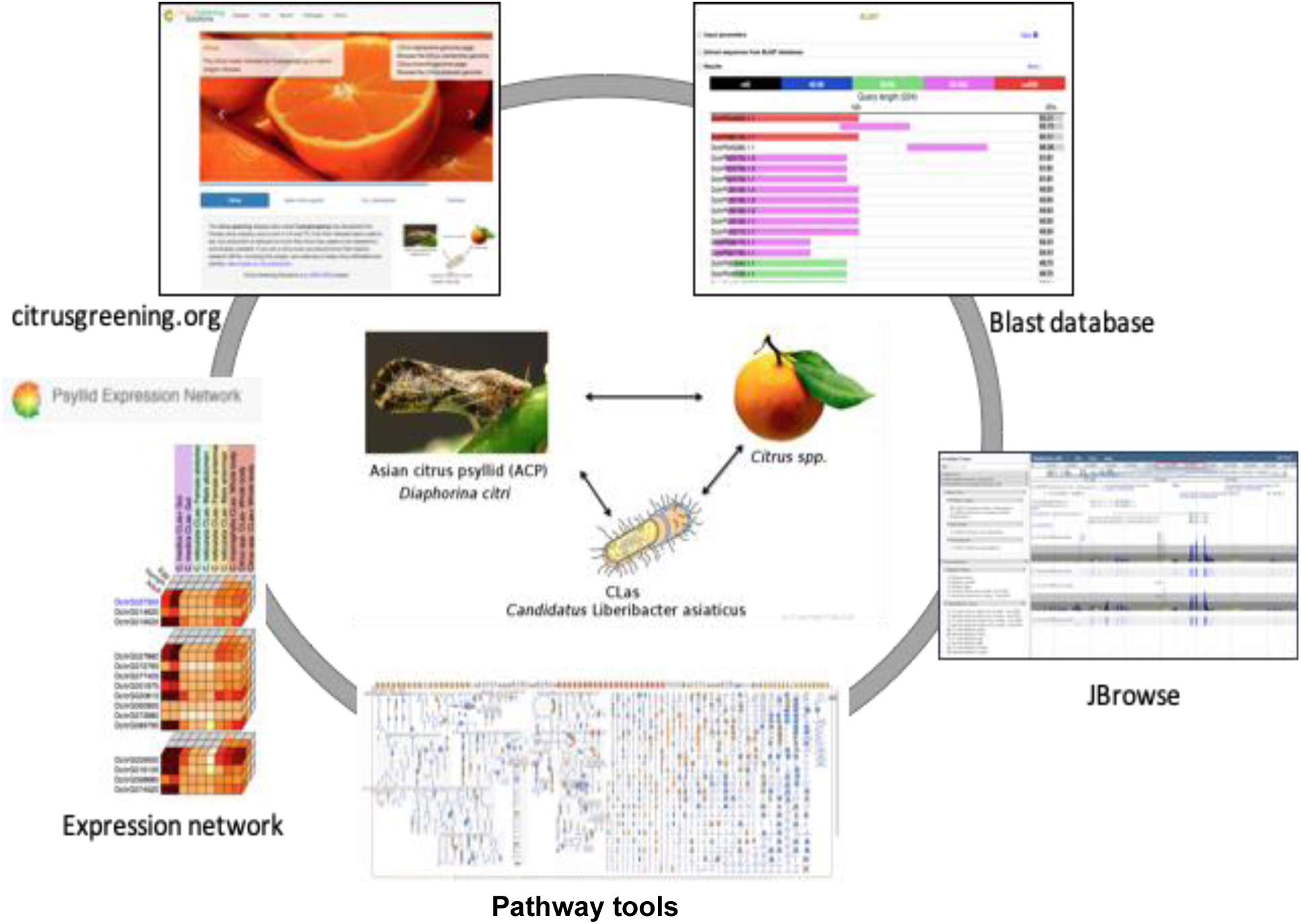
Citrusgreening.org, an open access web portal for the three components (host, vector and pathogen) of citrus greening disease.

**Figure 2:**
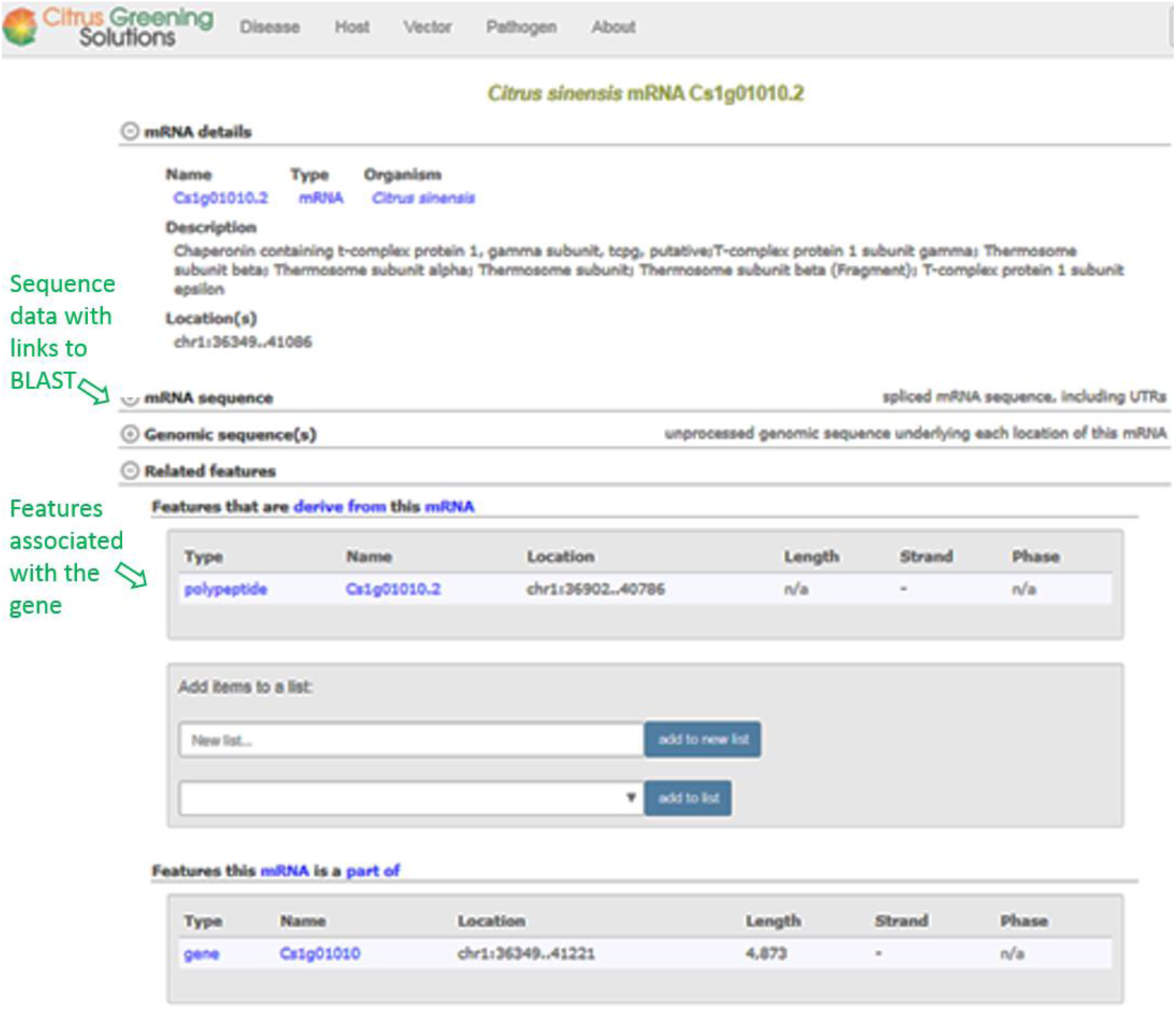
Gene feature page for *C. sinensis* with sub-sections with links to BLAST and component annotations.

Community driven curation is a central feature of Citrusgreening.org. Structural and functional curation of genes is implemented using the Apollo curation tool (Dunn et al. 2019). Gene annotators can curate gene models based on evidence from mapped orthologs as well as transcriptomics and *ab-initio* gene predictions. The manual curations from Apollo are periodically exported and merged with the automatically annotated gene models which are then released to the community (Saha et al. 2019). The platform has been used effectively for gene annotation by undergraduate students across multiple institutions and the curation workflow was described in Hosmani et al. (Hosmani et al. 2019). Besides Apollo, the Citrusgreening.org portal also has locus pages where additional information about a gene such as functional description, associated publications, GenBank sequences, images and ontology annotations can be added by approved locus editors. These features allow new information to be added incrementally over time and provide the community with a curated knowledge base to better understand organismal biology and interactions within the pathosystem.

### Citrus Expression Network (CEN) and Psyllid Expression Network (PEN)

Citrus Expression Network (CEN) and Psyllid Expression Network (PEN) are open-access, interactive and user friendly web tools to visualize gene expression profiles across multiple tissues, conditions and life stages. This tool is customized based on the Tomato Expression Atlas tool (Fernandez-Pozo et al. 2017) which is widely used by the Solanaceae research community. The expression network can be visualized with multiple views. Expression cube is the default view and shows expression of query gene and correlated genes in a heatmap (Figure 3). Expression values are also overlaid on a diagram of the organism to display tissue or stage specific activity in an experiment (Figure 4).

**Figure 3:**
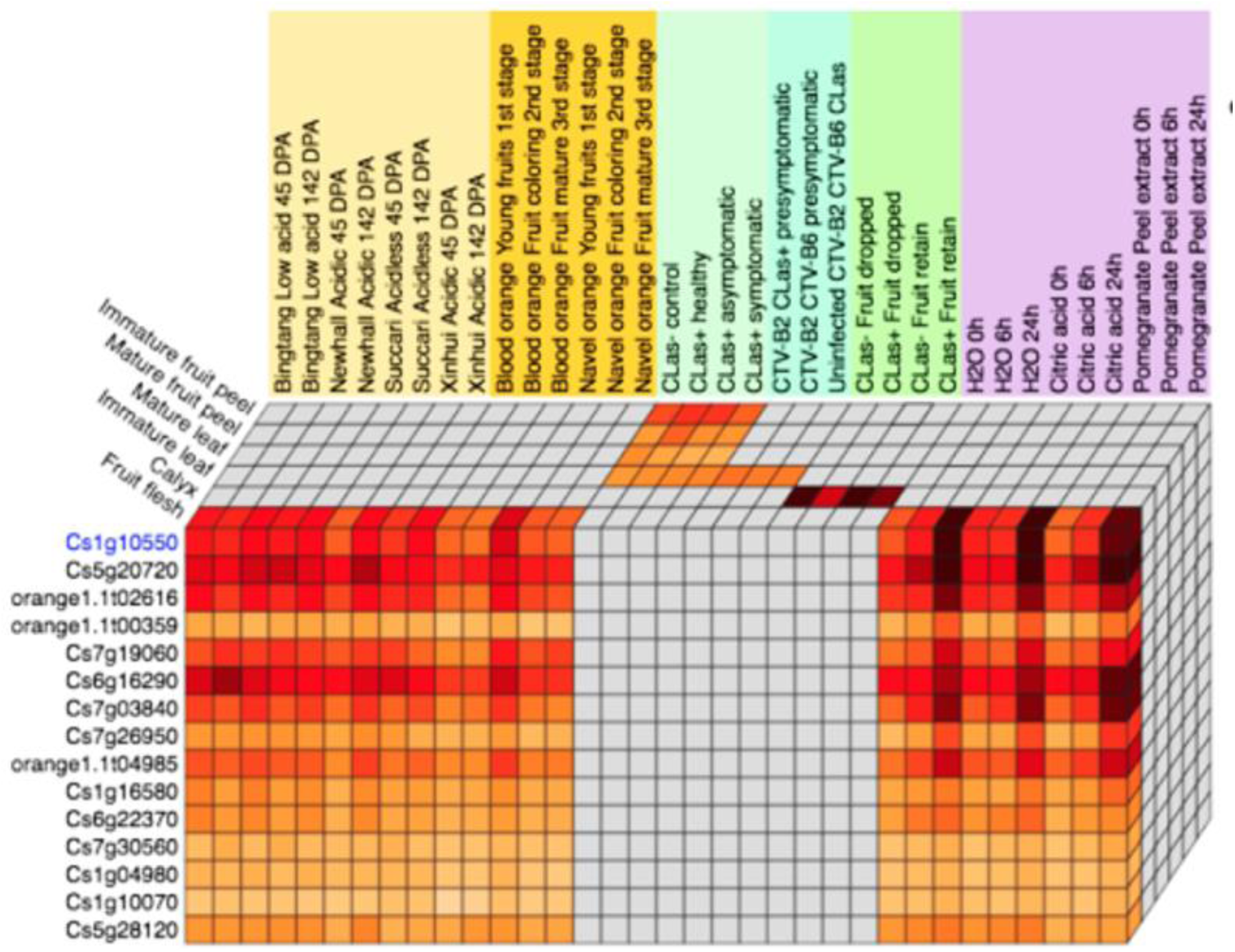
Expression cube from CEN. Heatmap displays expression values with the condition and citrus host on the X axis, genes on the Y axis and tissues on the Z axis. The genes are ranked by correlation and cells are colored by expression. Gray cells denote missing data.

**Figure 4:**
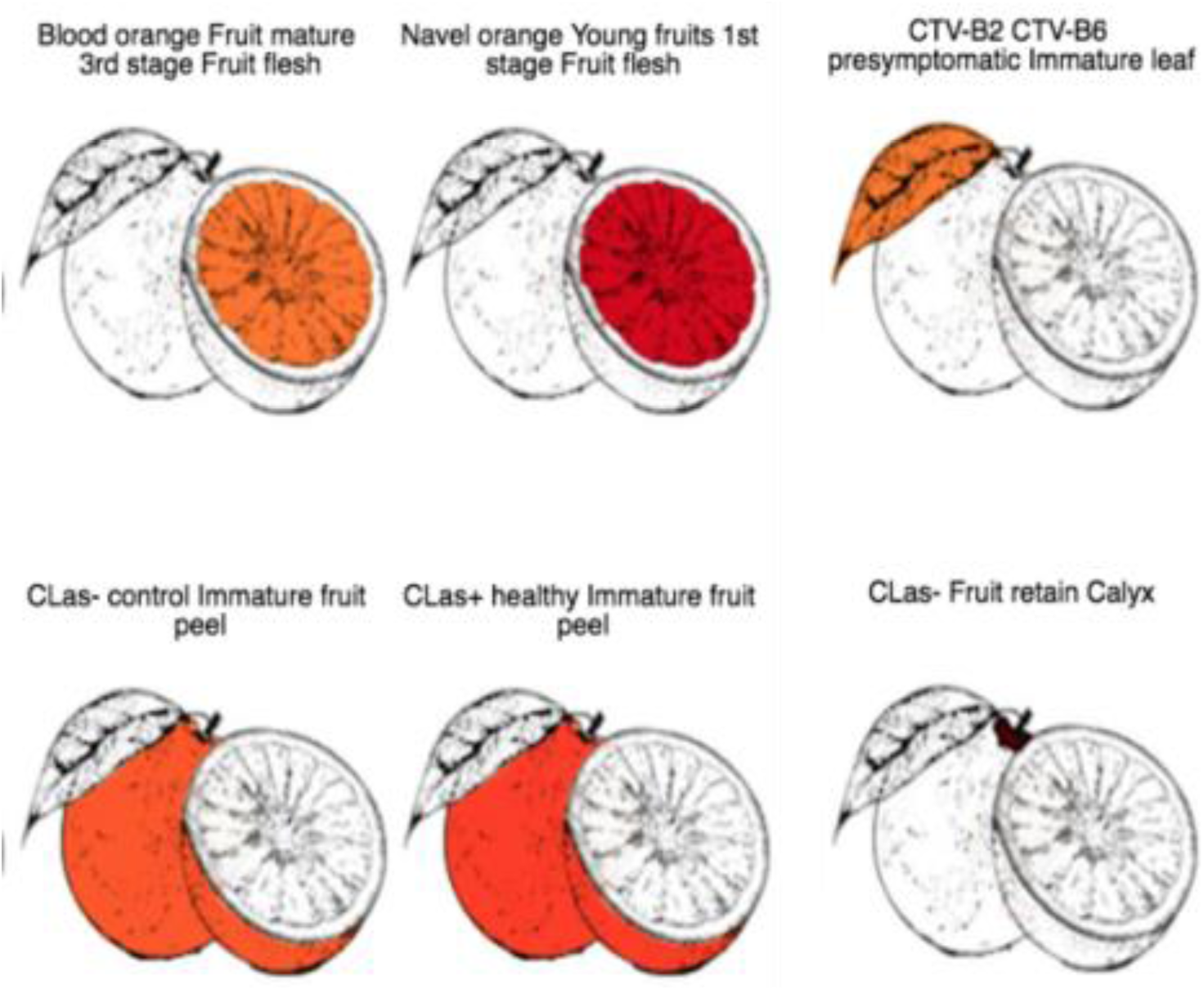
CEN expression images. Color coded images show gene expression according to tissue in the citrus fruit.

CEN contains transcriptomics data for *C. sinensis* and *C. clementina* from different citrus accesions (Bingtang, Newhall, Succari, Xinhui, Blood orange and Navel orange), stages (immature and mature fruit, young and mature leaf), tissues (fruit peel, fruit flesh, calyx and leaves), treatments (citric acid, peel extract) and infection states (CLas+, CLas-, CTV-B2, CTV-B6).

PEN, an expression atlas for psyllid combines transcriptomics and proteomics data from CLas infected and healthy individuals across multiple life stages (egg, nymph and adult), tissues (whole body, terminal abdomen, gut and antenna) and sexes (male and female). Insects were collected from a variety of hosts (*C. reticulata, C. macrophylla, C. sinensis, C. medica* and *C*.*clementina*). PEN contains data from multiple versions of the *D. citri* genome (Supplementary Table 2).

Gene transcription and translation patterns are critical for understanding how the underlying genome sequence translates into specific phenotypes at key developmental and infection stages. PEN and CEN allow visualization of the spatiotemporal context of gene expression and help elucidate function. This facilitates effective data analysis by enabling simultaneous visualization of correlated genes to develop novel hypotheses in addition to candidate gene identification. Detailed illustrations are used to show the spatial context of expressed genes (Figure 4). User can filter expression data on subsets of hosts, treatments, organs and developmental stages of interest. The resulting gene expression data can be downloaded from the expression cube panel as an Excel file for offline analysis.

The expression cube facilitates data analysis by enabling visualization of correlated genes, stages and conditions with color coded expression values. The cube can easily be broken down to see expression of specific or sets of correlated genes. Genes correlated with the query gene are sorted by correlation value on the Y axis. Expression images (Figure 4) reveal the internal anatomy of citrus overlaid with the expression profile in different tissues and stages in the fruit. Heatmap are used to show a traditional visualization of expression values in a condition by gene matrix. Additionally, scatterplots allow the user to identify outliers from selected samples.

### CitrusgreeningCyc pathway databases

Another important element of Citrusgreening.org is the identification and cataloguing of pathways in each organism involved in this disease complex. CitrusCyc (http://ptools.citrusgenomedb.org/) and DiaphorinaCyc (http://ptools-citrusgreening-ubuntu.sgn.cornell.edu/organism-summary?object=DIAPHORINA) are Pathway/Genome Databases (PGDB) that display metabolic pathways and metabolic maps for host and vector in the pathosystem. CitrusgreeningCyc (http://ptools-citrusgreening-ubuntu.sgn.cornell.edu/) also contains PGDBs for the pathogen CLas and other related Liberibacter species of interest for comparative analysis including Liberibacter cresens which is culturable and a model system for studying CLas (Table 1). The PGDBs have integrated tools for comparative analysis of pathways across multiple organisms and for analysis of user-supplied multi-omics data from metabolomics, transcriptomics and proteomics experiments. Users can also upload groups of genes or metabolites using the SmartTables feature (Karp et al. 2016) and perform multiple enrichment analysis. CitrusgreeningCyc was implemented with the pathway tools software suite (Karp et al. 2016) which was used to characterize metabolic pathways based on gene annotations and curated pathways in MetaCyc (Caspi et al. 2018). It currently contains 13 PGDBs with 2924 pathways, 21,732 enzymatic reactions, 37,895 enzymes and 16,465 compounds for 11 different species.

### Conclusions

The data and reports related to pathosystems are commonly focused on two or fewer organisms involved in plant disease complex. Moreover, this information is typically dispersed across a variety of journals and data repositories. There is a growing need for curating and connecting such resources so that the information is organized in standardized ontologies and is findable, accessible, interoperable and reusable (Sansone et al. 2019; Harper et al. 2018). Unified portals such as Citrusgreening.org can be used to provide a centralized data mining resource for a plant disease research community and stakeholders such as policy makers, farmers, extension agents, growers and regulatory authorities.

We have developed a data portal with a variety of comparative analysis tools in a flexible, extensible and community-curated system that integrates information from all the components of the tritrophic Huanglongbing or citrus greening disease complex. The next step is to extend this platform to incorporate ecological and climatic data in addition to plant phenotyping data from disease trials with related toxicology, insecticide resistance and behavioral assays.

While Citrusgreening.org has been created for use by molecular biologists and geneticists, our focus is to improve the usefulness of the site for citrus growers since they are valuable stakeholders who can help translate scientific advances into better integrated pest and disease management practices with subsequently higher yields in the grove. We have added information about tolerant citrus rootstocks to the site and will be expanding content relevant for growers. Integrated portals such as Citrusgreening.org will be a vital resource to tackle emerging diseases in agricultural systems and provide a platform to bring together and engage the stakeholders.

## Funding

All open-access fees and post-docs were funded through USDA-NIFA grant 2015-70016-23028.

## Methods

### Database implementation

The Citrusgreening.org platform is based on the Sol Genomics Network (solgenomics.net) architecture implemented primarily in the Perl language (Bombarely et al. 2011). The three-tiered data model consists of a user-facing view code, server side control and data modules with a relational database backend. The data is stored in a Chado relational schema (Mungall et al. 2007) with custom extensions in the PostgreSQL relational database system. A number of the site’s core functionality is based on the open source generic model organism database (GMOD) tools (http://gmod.org), such as Chado, Bio::Chado::Schema, Apollo and Jbrowse. The Blast services are provided by the NCBI BLAST+ suite [https://bmcbioinformatics.biomedcentral.com/articles/10.1186/1471-2105-10-421]. Flat files are used for a few storage purposes to complement the relational database, e.g. for storing large sequence data, metabolomics data and electrical penetration (EPG) graphs. All software source code and development logs are publicly viewable at http://github.com/solgenomics/sgn and https://github.com/solgenomics/citrusgreening.

### Pathways databases

Pathway/Genome Databases (PGDBs) are created using Pathway Tools (Karp et al. 2016) for pathway visualization and analysis. Organism specific pathway database was constructed using annotated proteins and curated pathways in MetaCyc (Caspi et al. 2018). In brief, PF files were created using genome annotation (GFF) and BLAST hits of proteins to sequences in Uniprot (UniProt Consortium 2019). Enzyme commission numbers were obtained for proteins from Uniprot database. GO terms (Harris et al. 2004) were obtained from InterProScan (Zdobnov and Apweiler 2001) and the Panther database.

Orthologous proteins were collected from *Drosophila megalonaster*, TrEMBL and Swiss-Prot insecta databases (Uniprot Taxonomy ID 50557) for the Psyllid and Araport 11, TrEMBL and Swiss-Prot plant databases (Uniprot Taxonomy ID 33090) for citrus. Sequence similarity was determined using BLAST (Altschul et al. 1990) with e-value 1e-5 and query coverage greater than 50%.

### Expression network

CEN contains public expression data for *Citrus sinensis* and *Citrus clementina* from RNA-Seq resources in the NCBI Sequence Read Archive (SRA) (Supplementary tables 4 and 5). PEN contains high-resolution expression data from proteomics and RNA-Seq datasets listed in Supplementary table 3. PEN was built using the *Diaphorina citri* v1.0 and v2.0 genome references. Gene expression values were obtained after mapping with HISAT2 (Kim et al. 2019) on the reference genome assembly. Expression counts were obtained using Stringtie and quantified using the transcripts per million reads (TPM) metric. Only genes with more than 1 TPM in at least one tissue were retained. All the expression values were provided to expression network (https://github.com/solgenomics/pen/, https://github.com/solgenomics/cen/) along with correlation matrix.

## Notes

https://citrusgreening.org/

