## Supplementary Data for "Citrusgreening.org: An open access and integrated systems biology portal for the Huanglongbing (HLB) disease complex"

Supp. Table 1. *Diaphorina citri* data

| **Data sets** |
|---|
| *Diaphorina citri* MCOT proteins |
| Diaphorina citri NCBI v100 proteins |
| *Diaphorina citri* genome v1.91 |
| *Diaphorina citri* genome v2.0 |
| *Diaphorina citri* OGS v1.0 proteins, transcripts, CDS, |
| *Diaphorina citri* OGS v2.0 proteins, transcripts, CDS, |
| *Diaphorina citri* Isoseq HQ |
| *Diaphorina citri de novo* transcriptome |

Supp. Table 2. PEN RNA-Seq datasets.

<table>
<tr><th>SRA</th><th>Data</th><th>Reference</th></tr>
<tr><td>SRR1259429</td><td>Citrus spp. CLas- Whole body Adult</td><td rowspan="4">https://journals.plos.org/plosone/article?id=10.1371/journal.pone.0130328</td></tr>
<tr><td>SRR1259432</td><td>Citrus spp. CLas+ Whole body Adult</td></tr>
<tr><td>SRR1259461</td><td>Citrus spp. CLas- Whole body Nymph</td></tr>
<tr><td>SRR1259434</td><td>Citrus spp. CLas+ Whole body Nymph</td></tr>
<tr><td>SRR2632316</td><td>C. reticulata CLas- Male antennae Adult</td><td rowspan="3">https://journals.plos.org/plosone/article?id=10.1371/journal.pone.0159372</td></tr>
<tr><td>SRR2632319</td><td>C. reticulata CLas- Female antennae Adult</td></tr>
<tr><td>SRR2632320</td><td>C. reticulata CLas- Male terminal abdomen Adult</td></tr>
</table>

| | | |
|---|---|---|
| SRR2632321 | *C. reticulata* CLas- Female terminal abdomen Adult | |
| SRR602249 | *C. macrophylla* CLas- Whole body Adult | https://www.jgenomics.com/v02p0054.htm#headingA5 |
| SRR610529 | *C. macrophylla* CLas- Whole body Nymph | |
| SRR610530 | *C. macrophylla* CLas- Whole body Egg | |
| SRR5514656 | ACP-Gut-Healthy-rep1a | https://journals.plos.org/plosone/article?id=10.1371/journal.pone.0179531#sec002 |
| SRR5514657 | ACP-Gut-Healthy-rep1b | |
| SRR5514655 | ACP-Gut-Clas-rep4b | |
| SRR5514654 | ACP-Gut-Clas-rep4a | |
| SRR5514651 | ACP-Gut-Clas-rep2b | |
| SRR5514649 | ACP-Gut-Healthy-rep4b | |
| SRR5514643 | ACP-Gut-Clas-rep1b | |
| SRR5514642 | ACP-Gut-Clas-rep1a | |
| SRR5514653 | ACP-Gut-Clas-rep3b | |
| SRR5514652 | ACP-Gut-Clas-rep3a | |
| SRR5514650 | ACP-Gut-Clas-rep2a | |
| SRR5514648 | ACP-Gut-Healthy-rep4a | |
| SRR5514647 | ACP-Gut-Healthy-rep3b | |
| SRR5514646 | ACP-Gut-Healthy-rep3a | |
| SRR5514645 | ACP-Gut-Healthy-rep2b | |
| SRR5514644 | ACP-Gut-Healthy-rep2a | |

Supp. Table 3. CEN *Citrus clementina* RNA-Seq data sets

| **SRA** | **Data** | **Reference** |
|---|---|---|
| SRR2828337 | Nour 3 DAF | PRJNA300206<br><br>https://www.ncbi.nlm.nih.gov/geo/query/acc.cgi?acc=GSE74384 |
| SRR2828338 | Nour 3 DAF | |
| SRR2828339 | Nour 3 DAF | |
| SRR2828340 | Nour 3 DAF | |
| SRR2828341 | Nour 3 DAF | |
| SRR2828342 | Nour 3 DAF | |
| SRR2828343 | Nour 3 DAF | |
| SRR2828344 | Nour 7 DAF | |
| SRR2828345 | Nour 7 DAF | |
| SRR2828346 | Nour 7 DAF | |
| SRR2828347 | Nour 7 DAF | |
| SRR2828348 | Nour 7 DAF | |
| SRR2828349 | Nour 7 DAF | |
| SRR2828350 | Nour 7 DAF | |
| ERR1307080 | Arrufatina_189_DPA | https://www.ncbi.nlm.nih.gov/bioproject/PRJEB12880/ |
| ERR1307081 | Clementina_126_DPA | |
| ERR1307082 | Hernandina_126_DPA | |

| | | |
|---|---|---|
| ERR1307083 | Hernandina_126_DPA | |
| ERR1307084 | Clementina_154_DPA | |
| ERR1307085 | Clementina_154_DPA | |
| ERR1307086 | Arrufatina_154_DPA | |
| ERR1307087 | Arrufatina_154_DPA | |
| ERR1307088 | Hernandina_154_DPA | |
| ERR1307089 | Hernandina_154_DPA | |
| ERR1307090 | Clementina_189_DPA | |
| ERR1307091 | Clementina_189_DPA | |
| ERR1307092 | Arrufatina_189_DPA | |
| ERR1307093 | Arrufatina_189_DPA | |
| ERR1307094 | Hernandina_189_DPA | |
| ERR1307095 | Hernandina_189_DPA | |
| ERR1307096 | Clementina_240_DPA | |
| ERR1307097 | Clementina_240_DPA | |
| ERR1307098 | Hernandina_240_DPA | |
| ERR1307099 | Hernandina_240_DPA | |
| ERR1307100 | Hernandina_275_DPA | |
| ERR1307101 | Hernandina_275_DPA | |
| Open flower | ERR768567 | https://www.ncbi.nlm.nih.gov/bioproject/PRJEB6342/ |

| | |
|---|---|
| Flowers (white button) | ERR776771 |
| Flowers (green button) | ERR776773 |
| Flowers (petals elongation) | ERR776774 |

Supp. Table 4. CEN *Citrus sinensis* RNA-Seq data sets

| **SRA ID** | **Data** | **Reference** |
|---|---|---|
| SRR3176495 | Succari 45 DPA | https://www.ncbi.nlm.nih.gov/bioproject/?term=PRJNA312406 |
| SRR3176496 | Succari 45 DPA | |
| SRR3176497 | Succari 45 DPA | |
| SRR3176498 | Bingtang 45 DPA | |
| SRR3176499 | Bingtang 45 DPA | |
| SRR3176500 | Bingtang 45 DPA | |
| SRR3176501 | Newhall 45 DPA | |
| SRR3176502 | Newhall 45 DPA | |
| SRR3176503 | Newhall 45 DPA | |
| SRR3176504 | Xinhui 45 DPA | |
| SRR3176505 | Xinhui 45 DPA | |
| SRR3176506 | Xinhui 45 DPA | |
| SRR3176507 | Succari 142 DPA | |
| SRR3176508 | Succari 142 DPA | |

| | | |
|---|---|---|
| SRR3176509 | Succari 142 DPA | |
| SRR3176510 | Bingtang 142 DPA | |
| SRR3176511 | Bingtang 142 DPA | |
| SRR3176512 | Bingtang 142 DPA | |
| SRR3176513 | Newhall 142 DPA | |
| SRR3176514 | Newhall 142 DPA | |
| SRR3176515 | Newhall 142 DPA | |
| SRR3176516 | Xinhui 142 DPA | |
| SRR3176517 | Xinhui 142 DPA | |
| SRR3176518 | Xinhui 142 DPA | |
| SRR5581496 | *Citrus sinensis* cv.zaohong blood orange fruit coloring at the 2nd stage | https://www.ncbi.nlm.nih.gov/bioproject/?term=PRJNA387319 |
| SRR5581497 | *Citrus sinensis* cv.zaohong blood orange fruit coloring at the 2nd stage | |
| SRR5581505 | *Citrus sinensis* cv.zaohong blood orange fruit coloring at the 2nd stage | |
| SRR5581498 | *Citrus sinensis* cv.zaohong blood orange young fruits at the 1st stage | |
| SRR5581499 | *Citrus sinensis* cv.zaohong blood orange young fruits at the 1st stage | |
| SRR5581500 | *Citrus sinensis* cv.zaohong blood orange young fruits at the 1st stage | |
| SRR5581501 | *Citrus sinensis* cv. 21st century navel orange young fruits at the 1st stage | |
| SRR5581502 | *Citrus sinensis* cv. 21st century navel orange young fruits at the 1st stage | |
| SRR5581503 | *Citrus sinensis* cv. 21st century navel orange young fruits at the 1st stage | |

| | | |
|---|---|---|
| SRR5581506 | *Citrus sinensis* cv. 21st century navel orange fruit mature at the 3rd stage | |
| SRR5581508 | *Citrus sinensis* cv. 21st century navel orange fruit mature at the 3rd stage | |
| SRR5581509 | *Citrus sinensis* cv. 21st century navel orange fruit mature at the 3rd stage | |
| SRR5581504 | *Citrus sinensis* cv. 21st century navel orange fruit coloring at the 2nd stage | |
| SRR5581510 | *Citrus sinensis* cv. 21st century navel orange fruit coloring at the 2nd stage | |
| SRR5581511 | *Citrus sinensis* cv. 21st century navel orange fruit coloring at the 2nd stage | |
| SRR5581507 | *Citrus sinensis* cv.zaohong blood orange fruit mature at the 3rd stage | |
| SRR5581512 | *Citrus sinensis* cv.zaohong blood orange fruit mature at the 3rd stage | |
| SRR5581513 | *Citrus sinensis* cv.zaohong blood orange fruit mature at the 3rd stage | |
| SRR5821806 | CLas+ fruit dropped | https://www.ncbi.nlm.nih.gov/bioproject/?term=PRJNA394061 |
| SRR5821807 | CLas+ fruit dropped | |
| SRR5821808 | CLas+ fruit dropped | |
| SRR5821809 | CLas+ fruit dropped | |
| SRR5821810 | CLas- fruit dropped | |
| SRR5821811 | CLas- fruit dropped | |
| SRR5821812 | CLas- fruit dropped | |
| SRR5821813 | CLas- fruit dropped | |
| SRR5821814 | CLas+ fruit retain | |
| SRR5821815 | CLas+ fruit retain | |

| | | |
|---|---|---|
| SRR5821816 | CLas+ fruit retain | |
| SRR5821817 | CLas+ fruit retain | |
| SRR5821818 | CLas- fruit retain | |
| SRR5821819 | CLas- fruit retain | |
| SRR5821820 | CLas- fruit retain | |
| SRR5821821 | CLas- fruit retain | |
| SRR867166 | Clas - control | https://www.ncbi.nlm.nih.gov/bioproject/?term=PRJNA428949 |
| SRR867394 | CLAS+ apparently healthy | |
| SRR867395 | CLAS+ asymptomatic | |
| SRR867396 | CLAS + symptomatic | |
| SRR867397 | Clas - control | |
| SRR867398 | CLAS+ apparently healthy | |
| SRR867421 | CLAS+ asymptomatic | |
| SRR867423 | CLAS + symptomatic | |
| SRR867425 | Clas - control | |
| SRR867426 | CLAS+ apparently healthy | |
| SRR867427 | CLAS+ asymptomatic | |
| SRR867431 | CLAS + symptomatic | |
| SRR867435 | Clas - control | |
| SRR867442 | CLAS+ apparently healthy | |

| | |
|---|---|
| SRR867446 | CLAS+ asymptomatic |
| SRR867449 | CLAS + symptomatic |
